# A novel integrated framework to identify and characterize regional-scale pest insect dispersal

**DOI:** 10.1101/2025.02.26.640127

**Authors:** F. Dargent, M.S. Reich, M. Miller, K. Studens, N. Benvidi, K. Perrault, J. Aibueku, B. Holmes, C.P. Bataille, J.N. Candau

## Abstract

Forest pest insects cause major socio-economic impacts, global losses of millions of dollars, and ecosystem changes. A key challenge for their management is tracing regional dispersal events critical to outbreak dynamics. We developed an integrated tracing framework for pest insects by combining isotope geolocation, ecological data, and atmospheric modeling, and applied this framework to the eastern spruce budworm moth (*Choristoneura fumiferana*), the most severe defoliator of the North American boreal forest, to trace outbreak dispersal events. We first generated a North American model of bioavailable sulfur isotope (*δ*^34^S) variation in space (isoscape), which predominantly varied in response to oceanic sulfate deposition, and then calibrated it to spruce budworm tissues of known origin. We used an automated trap network with high temporal resolution to collect samples and identify potential immigration events of eastern spruce budworm to Nova Scotia, Canada. We traced the natal origin of these immigrants by integrating high-probability regions derived from *δ*^34^S probabilistic assignments and HYSPLIT atmospheric dispersal models. Since high larval density is a strong predictor of budworm defoliation and emigration, HYSPLIT atmospheric dispersal models, which integrated spruce budworm behavioral constraints (e.g., flight velocity, altitude, and temperature thresholds), were started from defoliated areas to narrow-down the area of natal origin and estimate the migration route. We find that this integrated framework allows to narrow down the region of pest origins, restricting it to a few possible locations and demonstrating long-distance dispersal of spruce budworm across ∼400Km over the Gulf of St. Lawrence. Our framework demonstrates the utility of *δ*^34^S geolocation in insect tracing, and that combining isotopic data with ecological indicators and atmospheric modeling offers an unprecedented resolution in understanding insect dispersal ecology. The approach is transferable to trace other migratory insect species to address conservation, agriculture, and bio-surveillance needs in the context of global environmental change.

## 1. Introduction

Insect pests are a major economic and societal challenge (Bradshaw et al. 2016). Each year they cost the global economy hundreds of billions of dollars in crop losses (Culliney 2014, FAO 2021), with outbreaks leading to further complex impacts on ecosystem services via changes to ecosystem structure and function. These costs are expected to increase in the future as human populations and demand for agricultural and forestry products continue to grow, climate change exacerbates the frequency and extent of forest insect outbreaks (Jactel et al. 2019) and crop losses (Deutsch et al. 2018), and changing wind patterns facilitate pest dispersal to new regions (Harvey et al. 2023). The scale and magnitude of these impacts is closely tied to each species’ dispersal capacity because it influences the rate of pest spread, their recurrence, and the likelihood of establishment (Heimpel and Asplen 2011, Asplen 2018).

Successful management strategies to forecast and mitigate pest risks to ecosystems, agriculture and human societies requires, first, developing effective tools to evaluate dispersal (Coulson et al. 1993). Such tools should provide accurate spatio-temporal knowledge of the source, trajectory, and endpoint of pest movements, which allows the identification of causal mechanisms that trigger insect movement, and the ecological consequences at the landing sites. Yet, several factors have limited our ability to identify and characterize insect dispersal events. First, the vast magnitude and frequency of insect dispersal has only been demonstrated recently (Satterfield et al. 2020), and incorporates a broad diversity of movement strategies, ranging from active to passive transport, stochastic to defined routes, single-generation to multigenerational processes, involving individual or synchronized behaviors, and local to continental scales (Chapman et al. 2015, Satterfield et al. 2020). Second, typical animal tracking methods (e.g., radio telemetry, tagging) are challenging to implement on insects due to low body mass, while mark and recapture is hampered by their short lifespan, large populations, and often stochastic dispersal routes (El Sheikha 2019). Third, while more recent tracing tools, like radar and population genetics, show promise, their successful implementation can be highly context-specific. For example, radar technology needs extensive infrastructure and is limited to detecting massive migration events (e.g., Shamoun-Baranes et al. 2014), and migration and outbreaks often increase gene flow and erase the genetic population structure that is necessary for population genetic tools (e.g., Lumley et al. 2020). Isotopic tracers that are naturally incorporated in insect tissues coupled with atmospheric dispersal models based on accurate capture or take-off times, could potentially circumvent many of these limitations.

Isotopes have been used for decades to reconstruct long-distance insect migration because they represent intrinsic markers of geographic origin (Quinby et al. 2020). While some isotope tracers are commonly used (e.g., hydrogen and strontium isotopes; Reich et al. 2021), the development of novel geolocators from isotopes of elements that vary independently from these would further revolutionize the possibility to track insect migration at high resolution. Sulfur isotopes (referred to as *δ*^34^S) vary on the landscape with atmospheric deposition processes or geology, and have shown promising potential for geolocation in archeology (Bataille et al. 2021) and bird ecology (Brlík et al. 2023). These isotopes vary predictably on the landscape and when insect larvae develop at a given location, the tissues metabolized at that site inherit a local isotope “fingerprint” (Quinby et al. 2020). If those tissues have limited metabolic recycling (e.g., wings) (Lindroos et al. 2023) and if the studied isotope is not easily contaminated (Reich et al. 2023), the tissue can preserve that isotopic fingerprint reflecting the site of natal origin. By comparing the isotopes of a migratory individual to a reference map predicting isotopes on the landscape (i.e., isoscape), one can estimate the origin of an individual. Newton (2021) demonstrated that moths collected across the UK showed consistent spatial patterns with *δ*^34^S gradient from the coast to inland. However, *δ*^34^S has not been tested as a geolocator for insect migration. Sulfur isotopes are attractive as an insect geolocator because sulfur is a macronutrient present in measurable amounts in many insect tissues through two protein-forming amino acids, the essential methionine and the non-essential cysteine (Tcherkez and Tea 2013). The sulfur isotope cycle is also well-characterized; primary producers (i.e., plants) obtain sulfur primarily from soils and more rarely through atmospheric uptake as sulfur oxides and carbonyl sulfides (Tcherkez and Tea 2013). In most ecosystems, soil sulfur is derived from atmospheric deposition of marine sulfates, which have a homogeneous and elevated *δ*^34^S (+21 ± 0.2 ‰), deposited either by short-distance dry deposition of sea spray or wet deposition of sulfates dissolved in precipitation (Supplementary Figure 1). Conversely, geological influences, microbial activity and other atmospheric deposition processes dominate sulfur cycling in ecosystems with low oceanic sulfate deposition (Nehlich 2015). The complex cycling and multiple isotopically-distinct sources of sulfur in ecosystems might lead to isotope patterns independent from hydrogen and strontium providing a new tool for high-resolution insect tracing.

Similarly, atmospheric particle dispersal models have improved the tracing and forecast of wind-assisted insect dispersal (NOAA 2020). These models use time-dependent gridded meteorological data to calculate transport and diffusion processes of air parcels in the atmosphere, and can be applied to simulate the release, transport, and deposition of particles, including wind-transported insects, on a given timeframe and obtain their density fields, mean trajectories, and deposition rates (e.g., Stein et al. 2015). This process’ precision can be further enhanced by incorporating insect behavioral and ecological parameters such as flight phenology, biophysical constraints, mass-dependent effects, and population dynamics (e.g., Otuka et al. 2023).

However, to be effective for investigating insect dispersal, atmospheric particle dispersal models require that the location and time of the insect’s arrival be known. Remote monitoring systems, such as automated trap networks, provide a temporal context (e.g., time of capture) for individual specimens that can be used to constrain the spatiotemporal range of potential atmospheric particle dispersal simulations (Preti et al. 2021). Coupling samples collected from trap networks with isotope geolocation and atmospheric particle dispersal model should offer significant promise for enhancing insect pest management and improving decision support systems.

In this study, we use the eastern spruce budworm moth (*Choristoneura fumiferana* Clemens, hereafter spruce budworm) (Lepidoptera: Tortricidae) as a model species to evaluate the effectiveness of combining sulfur isotopes, atmospheric modeling, and ecological inferences to trace migratory insects with a focus on improving insect pest management strategies. The spruce budworm is an excellent study species as it is the most severe and pervasive native defoliator of conifers in the North American boreal forest (Morris 1963, MacLean 2019), causing substantial ecological change (e.g., Bouchard et al. 2006, Eveleigh et al. 2007) and major negative economic and social impacts when outbreaks occur (e.g., Liu et al. 2019). During outbreaks, spruce budworm population density increases several orders of magnitude, leading to heavy defoliation of their spruce (*Picea* sp.) and balsam fir (*Abies balsamea*) hosts. The decrease in resource availability increases competition and the propensity of adult male moths and egg-carrying females to initiate mass exodus from outbreak areas (Morris 1963, Régnière and Nealis 2007). Once aloft over the tree canopy, millions of moths can be carried by the wind over long distances (up to 450 km) (Greenbank et al. 1980), and this mass migration of budworm moths is thought to be the main driver of outbreak expansion in newly colonized areas (Morris 1963).

Efficient management of these outbreaks requires early warning systems capable of forecasting the immigration of moths to target areas vulnerable to future outbreaks, combined with preventive measures that anticipate and stop outbreak spread (Johns et al. 2019). Yet, limited capacity to trace spruce budworm migration at a high temporal and spatial resolution, remains an important challenge to evaluating the biotic and abiotic drivers of take-off (emigration) and landing (immigration) that contribute to outbreaks. As a consequence, the current approach to monitoring, controlling and mitigating the growing spruce budworm outbreak requires in-situ monitoring of vast forest areas, which is expensive and time-consuming (Natural Resources Canada 2021) and would greatly benefit from developing novel integrated geolocation tools for insect dispersal.

To evaluate the use of an integrated approach for insect geolocation, we first developed a spruce budworm isoscape across eastern North America by 1) collecting and analyzing *δ*^34^S values in plants from across eastern North America, 2) developing a foliar *δ*^34^S (*δ*^34^S_foliar_) isoscape across the study area using random forest regression and geospatial remote sensing datasets and 3) conducting a rearing assay in the wild to relate the isotopic composition of adult moths with that of their host trees. We also tested for sources of intra-site variance in *δ*^34^S values, including tree species and needle age. We then leveraged automated pheromone traps across eastern Canada, to collect immigrant moths, classified as such from independent sources of evidence including phenology, capture rates, and other isotopes. We analyzed their *δ*^34^S values and used isotope-based geographic assignments along with field-derived defoliation maps (indicative of budworm activity and migration propensity) to determine their most likely region of natal origin. We simulated potential dispersal pathways from defoliated regions and selected those that reached the capture location of these immigrants, and combined them with the most likely sites of natal origin to refine our origin predictions. Finally, we discuss the broader impacts of this integrated framework as a tool to inform insect pest management.

## 2. Methods

### 2.1. Foliar Sulfur Isoscape

#### 2.1.1. Foliar sample collection

Since plants are the primary source of sulfur for folivore insect larvae, we expect a predictable relationship between local *δ*^34^S_foliar_ and the *δ*^34^S of adult insects (*δ*^34^S_moth_) that fed on those plants as larvae. Thus, a *δ*^34^S_moth_ isoscape could be readily constructed from a *δ*^34^S_foliar_ isoscape (Hobson 2019). To develop the *δ*^34^S_foliar_ isoscape for North America, we collected mature needles from the host plants of spruce budworm, balsam fir (*Abies balsamea*) and spruce (*Picea sp.*), across, mostly, eastern Canada, from sites encompassing a broad range of environmental, geological, geographic, anthropogenic and climatic conditions. These samples were collected between 2019 and 2022 by collaborators and volunteers at 157 sites, and their latitude and longitude were recorded. At 77 of these sites, we collected branches at 1.5 meters above ground from three trees less than 10 meters apart. At the remaining 80 sites, only one tree was sampled. In the laboratory, each branch was subsampled and, when possible, species was recorded before the samples were stored in kraft envelopes in a dry room. To cover a broader geographic range, we also analyzed *δ*^34^S_foliar_ in common milkweed (*Asclepias syriaca*) collected from 39 sites across the United States in the summer of 2018 (see Reich et al. 2021). In total, samples were collected from 197 sites (Pinaceae n=157, milkweed n=39) between 2018 and 2023 (most milkweed samples were collected in 2018).

We tested the possible intra-site δ*^3^*^4^S_foliar_ variation caused by tree species differences by collecting and comparing spruce and balsam fir needles from the same sites (n=8). Additionally, since spruce budworm prefers feeding on more-nutritious immature (i.e., young) foliage (Fuentealba and Bauce 2012), we collected both young needles and fully mature needles from the same trees at seven sites, and compared their isotopic composition.

#### 2.1.2. Foliar sample preparation and isotope analysis

Dry needles were aggregated by site, rinsed twice with deionized water to remove surficial particles, and dried at 50 °C for 48h. Milkweed leaves were aggregated by site and rinsed with deionized water in an ultrasonic bath and dried at 70 °C for seven days. After drying, dry needles and leaves were ground to a fine powder and stored in glassine envelopes.

Sulfur isotopes were analyzed at the Jan Veizer Stable Isotope Laboratory at the University of Ottawa. We micro-weighed a target mass of 60 ± 6 mg of conifer needle powder (S%= 0.09 ± 0.03), and 25 ± 2.5 mg of milkweed leaf powder (S%= 0.51 ± 0.18), into tin capsules. Samples were combusted through an Elemental Analyser Isotope Cube in S mode (Elementar, Germany) with combusted SO_2_ from the sample carried to a Delta Plus XP IRMS (ThermoFinnigan, Germany) equipped with a ConFlo IV. The ^34^S/^32^S ratios were obtained from *m/z* 48, 49 and 50 of SO^+^ produced from SO_2_. Standards used for three-point calibration were sulfide-based IAEA-S1 (argentite, δ*^34^S* = -0.3 ‰ ± 0.03), IAEA-S2 (argentite, *δ*^34^S = 22.7 ‰ ± 0.08) and IAEA-S3 (argentite, *δ*^34^S = -32.6 ‰ ± 0.08) international standards (International Atomic Energy Agency 2020; IAEA, Vienna, Austria); AG-2 (argentite, *δ*^34^S = -0.6 ‰) was used as an internal blind standard. Sulfur isotope values are reported to the international scale Vienna Canyon Diablo Troilite (VCDT). The measured values for AG-2 among runs varied between -0.4‰ and -1.1‰ (among run average = -0.8± 0.3‰, n = 9), and were within the laboratory long-term reported value of -0.6‰ and uncertainty of 0.3‰. Analytical precision of these measurements is based on the reproducibility of 22 foliar powder samples which had a mean difference between duplicates of 0.16 ‰ ± 0.15‰.

#### 2.1.3. Foliar isoscape modeling

We first built a *δ*^34^S_foliar_ isoscape using random forest regression following the methods of Bataille et al. (2020, 2021) and Reich et al. (2024). The model incorporated the newly produced δ*^34^*S data from foliar samples across eastern North America and continuous geospatial variables representing geological, atmospheric, climatic or environmental controls of sulfur isotope cycling. The assembled covariates have been previously described (Reich et al. 2024 and Supplementary Table 1). The tuning and optimization of the random forest model has been described in previous work (Bataille et al. 2020). Briefly we used the “tuneRF” function of the *caret* package (Kuhn 2008) using RMSE (root mean square error) through a 10-fold cross-validation approach with five repetitions where 80% of the data was used for training at each iteration. We used the *VSURF* package (Genuer et al. 2015) to maximize model performance while minimizing the number of predictors. Partial dependence plots were also used to evaluate the relationship between *δ*^34^S_foliar_ values and the selected predictors. We estimated model uncertainty using quantile random forest to calculate a 68.27% prediction interval (Funck et al. 2021) which we used to then estimate the standard deviation (σ) of predictions at each pixel with: a ≈ q(0.841) - q(0.159).

### 2.2. Spruce budworm moth-calibrated *δ*^34^S isoscape

#### 2.2.1. Known-origin moths sample collection

The relationship between *δ*^34^S_foliar_ and *δ*^34^S_moth_ values for spruce budworm is unknown, although previous studies have shown limited overall fractionation between resource and tissue in invertebrates (mean Δ^34^S = +0.6 ±0.3‰, non-significantly different from 0) and a slight depletion of ^34^S relative to the food source in other animals (Raoult et al. 2024). As spruce budworm ingest plants and liquid only during the larval stages and have a short life span (∼10 days) as adults (Miller 1963), the isotopic composition of adult tissues likely reflects that of the larval diet.

To assess the relationship between *δ*^34^S_foliar_ and *δ*^34^S_moth_ values and account for potential trophic level isotopic fractionation we collected spruce budworm on up to three adjacent trees at 19 sites in Ontario, six sites in Newfoundland, and three sites in New Brunswick (“known-origin” samples). Following Contina et al. (2022), the sites were selected to include extreme values and a broad range of environmental conditions known to influence sulfur isotope cycling (e.g. Bataille et al. 2021, Brlík et al. 2023). We favoured collecting field samples over rearing spruce budworm on isotopically distinct diets to avoid potential bias in how sulfur in diet is incorporated in the tissues and better account for the potential incorporation of atmospheric deposition on ingested needles. For each known-origin site (except those in New Brunswick foe which we had no foliar samples), we collected foliar samples from the same trees that we sampled for spruce budworm, or within 10 meters of budworm collection (NL), and pooled them for foliar analysis as describe in section 2.1.1. To ensure they were true locals, most known-origin moths in our dataset were collected in the field as pupae and allowed to eclose in captivity, in paper bags at roughly 22°C (laboratory temperature). We recorded the geographic coordinates, collection date, site description, and match between plant and moth samples. To assess intra-site *δ*^34^S_moth_ variations, we aimed to collect 3-5 adults per site, but this was not possible at all sites, due to low density of budworm, failure to eclose, and attacks by Hymenoptera parasitoids.

#### 2.2.2. Moth sample preparation and isotope analysis

Each moth was cleaned in a 2:1v/v chloroform:methanol solution in three successive washes for 30 min, 3 h, and 15 min to remove lipids, dust, contaminants, and glue (i.e., for moths captured in automated traps). The samples were then air dried in a class-100 fume hood and stored in glassine envelopes. Insect wings are often preferred for isotope analysis because they have limited tissue turnover (Lindroos et al. 2023). However, lepidoptera wings have very low sulfur content (<0.01% in spruce budworm moths) as they are primarily composed of chitin (C_8_H_13_O_5_N)_n_ (Kaya et al. 2015). We analyzed sulfur isotopes using similar procedures as foliar samples (see Section 2.1), but smaller sample weights (2 ± 0.2 mg) with thoraces, and heads sampled in priority and part of the abdomen incorporated for low mass samples. Since spruce budworm are not known to eat or drink as adults in the wild, and are capital breeders (Rhainds 2015), it is likely that individual *δ*^34^S values of the adult reflect that of the larvae in most tissues. As moth tissues are made of chitin, we also regularly analyzed an internal chitin standard that we created from ground and homogenized spongy moth (*Lymantria dispar*, Linnaeus, 1758) heads and thoraces. This quality check standard gave δ*^34^S* = 2.7 ± 0.3 ‰ (n=3); those values support an analytical precision of ±0.3‰.

#### 2.2.3. Isoscape calibration to spruce budworm moth tissue

We generated a *δ*^34^S_moth_ isoscape using the *assignR* package (Ma et al. 2020). We used the *calRaster* function to fit a linear model relating *δ*^34^S_moth_ from the known-origin dataset and the random forest *δ*^34^S_foliar_ isoscape. To evaluate the quality of the calibrated isoscape, we used the *QA* function.

### 2.3. Automated trap collections and moth migration assessment

#### 2.3.1 Trap collections and inferences of migration

We used automated pheromone traps (Trapview+ model, EFOS d.o.o., Slovenia) deployed by the Canadian Forest Service and Provincial Governments to collect male spruce budworm across their distribution and expansion range (see Dargent et al. 2023 for details). These traps use sticky paper to collect specimens and take photos four times daily (at approximately 05:00, 11:00, 19:00 and 23:00) to determine the date and approximate time of capture of each individual. Using a combination of phenology modeling, climate conditions at the trap location, abundance of capture from previous days, and time of capture (Sanders et al. 1978, Greenbank et al. 1980), moths are qualitatively classified between locals and immigrants. Broadly, individuals captured in the evening and early night are generally considered local, whereas moths captured in the early morning or at a large and sudden peak in capture numbers reflect a potential immigration event. We selected specimens from two automated traps in Arisaig (45.75°N, −62.16°W) and Inverness (46.196°N, −61.294°W) in Nova Scotia from a known immigration event (i.e., July, 23rd 2020) to test and develop the sulfur isotope geolocation tool. The immigration event at these sites had been documented through several independent lines of evidences including: 1) a strong decline in local moth activity in the area followed by a sudden abundance of captures three to eight times higher than previous peak captures and 2) combined δ2H values and 87Sr/86Sr ratios from those captured individuals different from those of identified locals from previous weeks (Dargent et al. 2023).

### 2.4 Assignment of immigrant provenance

#### 2.4.1 Continuous-surface probabilistic assignment

We used the *pdRaste*r function in the *assignR* package (Ma et al. 2020) to generate continuous-surface probabilistic maps of natal origin for all specimens sampled at capture sites between 2019 and 2020, including purported locals and immigrants.

#### 2.4.2. HYSPLIT atmospheric dispersal trajectories

Independently, we tested whether the moths identified as potential immigrants on the night of July 22, 2020 in Arisaig and Inverness had potential wind-assisted trajectories corresponding to areas of high population density. As a reference for population density, we used defoliation maps from 2020 generated by the Ministère des Ressources Naturelles et des Forêts on the north shore of the Saint Lawrence River and the Gaspe peninsula (Ministère des Forêts 2020). We rasterized and resampled this map into a 1 km grid. New Brunswick, Newfoundland, and Nova Scotia annual surveys did not report evidence of spruce budworm outbreaks in 2020.

To evaluate the potential flight trajectories for the immigrants, we performed dispersal simulations using HYSPLIT (v. 5.2.0) from all defoliated cells. HYSPLIT uses the Lagrangian method to simulate transport and diffusion processes in air parcels (Stein et al. 2015). Following Greenbank et al. (1980), these dispersal simulations incorporated a “moth velocity” parameter (i.e., 2.5 m/s) (e.g., Otuka et al. 2023), that accounts for active flying by moths. We generated plausible flight trajectories for the dates of 2020-07-21 and 2020-07-22, starting at 19:00, 21:00, 23:00 ADT, at 300, 600, and 900 m altitude for a duration of 9 hours. We used meteorological data from the North American Mesoscale Forecast System at 12 km cells. We identified trajectories that landed inside or intersected a radius of 10 km around each trap, and evaluated temperature and altitude during the migratory flight trajectory to confirm that trajectories did not occur below the temperature flight threshold of spruce budworm (i.e., <15 °C) or got close to ground level (i.e., <250 m) which could have triggered landing before reaching the traps.

### 2.5 Statistical analyses

All statistical analyses were performed using R version 4.3.1 (R Core Team 2020, http://www.r-project.org). R scripts and data are available at OSF [*on acceptance*].

## 3. Results

### 3.1. Foliar sulfur isoscape

The *δ*^34^S of the plant samples ranged from -8.5 ‰ to 18.6 ‰ (mean = 7.46 ‰) and displayed a bimodal distribution (Figure 1A). The random forest model identified six predictors of *δ*^34^S_foliar_ in North America including sea salt aerosol, distance to the coast, average mineral dust deposition, wind speed, aridity index (i.e., mean annual precipitation divided by mean annual evapotranspiration), and potential evapotranspiration (Figure 1). These variables capture the known influences of atmospheric deposition, climate and dust inputs on sulfur isotopes cycling and *δ*^34^S spatial variation. The model explained 85.7% of the variance in *δ*^34^S values and had a root mean square error (RMSE) of 2.1 ‰, which is 7.6 % of the range of values we collected (Figure 1A, 1B). The residuals of our model were normally distributed but tended to overpredict lower *δ*^34^S_foliar_ values, particularly for samples collected in the Northwest Territories (Figure 1B).

**Figure 1:**
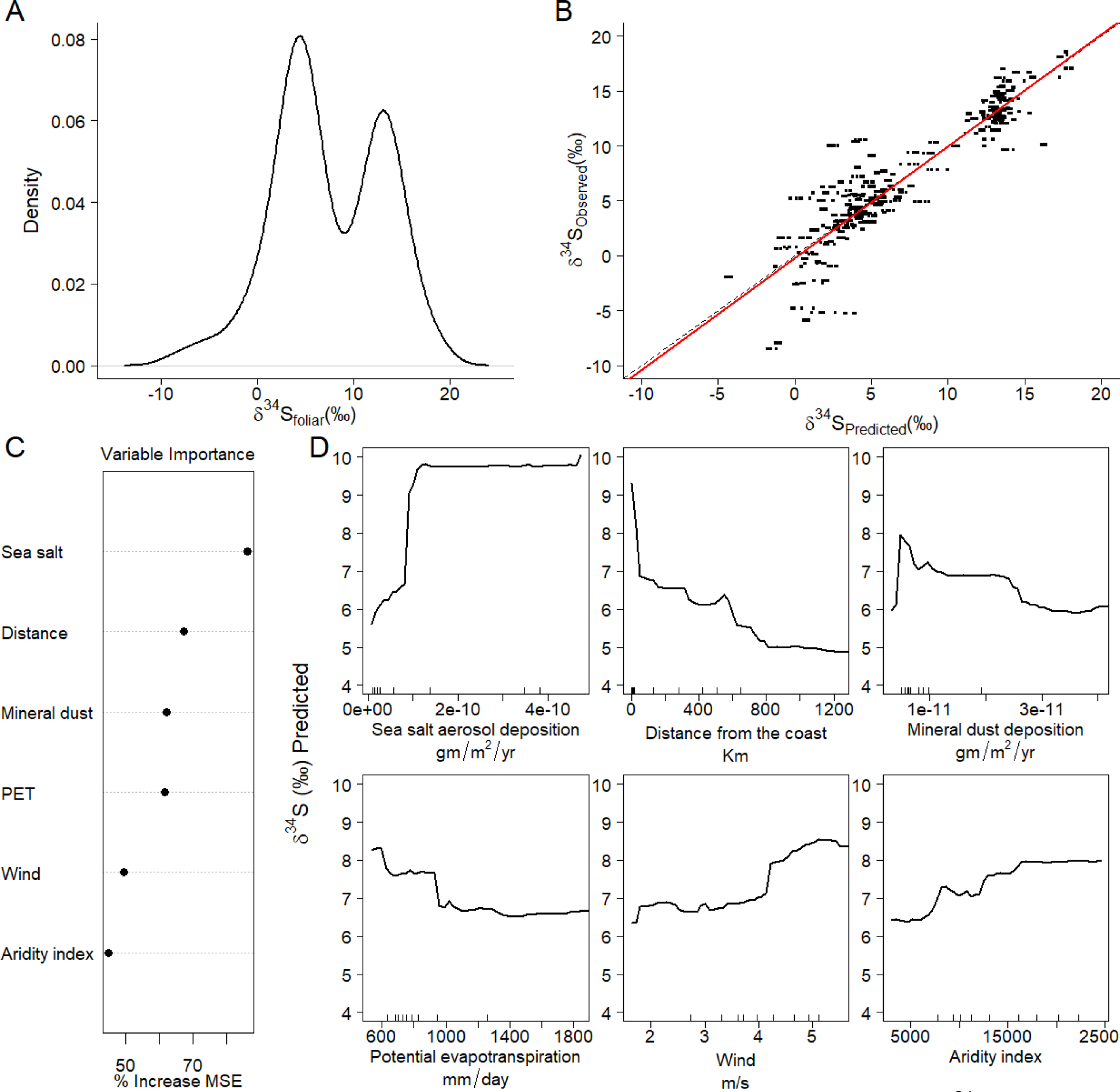
Analysis of foliar sulfur isoscape of North America. (A) Distribution of *δ*^34^S_foliar_ values in spruce *spp.*, balsam fir, and milkweed samples across North America. (B) Cross-validation of predicted vs. observed *δ*^34^S_foliar_ values using random forest regression. The red line represents the best-fit linear model, broken line a 1:1 relationship between predicted and observed values, R^2^ and Root Mean Squared Error (i.e., RMSE) of the model are reported. (C) Variable importance plots of *VSURF*-selected predictors based on increase in node purity. (D) Partial dependence plots of *VSURF*-selected predictors with predicted *δ*^34^S_foliar_ values and the predictors.

Among the significant predictors, salt aerosol deposition, a proxy of oceanic sulfate deposition, showed a threshold relationship with *δ*^34^S_foliar_ values with higher values near the coast, particularly in eastern Canada (Supplementary Figure 2A). Similarly, distance from the coast, aridity index and potential evapotranspiration showed lower *δ*^34^S_foliar_ values away from the coast or in dryer conditions. Wind velocity, which controls the distance and size of particles carried via atmospheric deposition (e.g., dry sea salt spray), showed a strong relationship with *δ*^34^S_foliar_ values, with higher *δ*^34^S_foliar_ within a narrow coastal band (Supplementary Figure 2B).

Dust aerosol deposition also showed a threshold relationship with *δ*^34^S_foliar_ values, following a broad latitudinal gradient increasing northward (Supplementary Figure 2C, 2D).

Overall, our *δ*^34^S_foliar_ isoscape reveals a strong gradient of decreasing *δ*^34^S_foliar_ with increasing distance from coastal areas, and very high *δ*^34^S_foliar_ values within a narrow coastal band influenced by oceanic winds. Away from the coast, *δ*^34^S_foliar_ values become more influenced by mineral dust deposition (Figure 2). Interestingly, despite the known influences of certain lithologies (e.g., black shales and evaporites) and anthropogenic pollution on ecosystem’s *δ*^34^S values (e.g., Nehlich 2015, Kabalika et al. 2020), the random forest model did not detect any direct influence of geological variables or human-derived atmospheric deposition variables.

**Figure 2:**
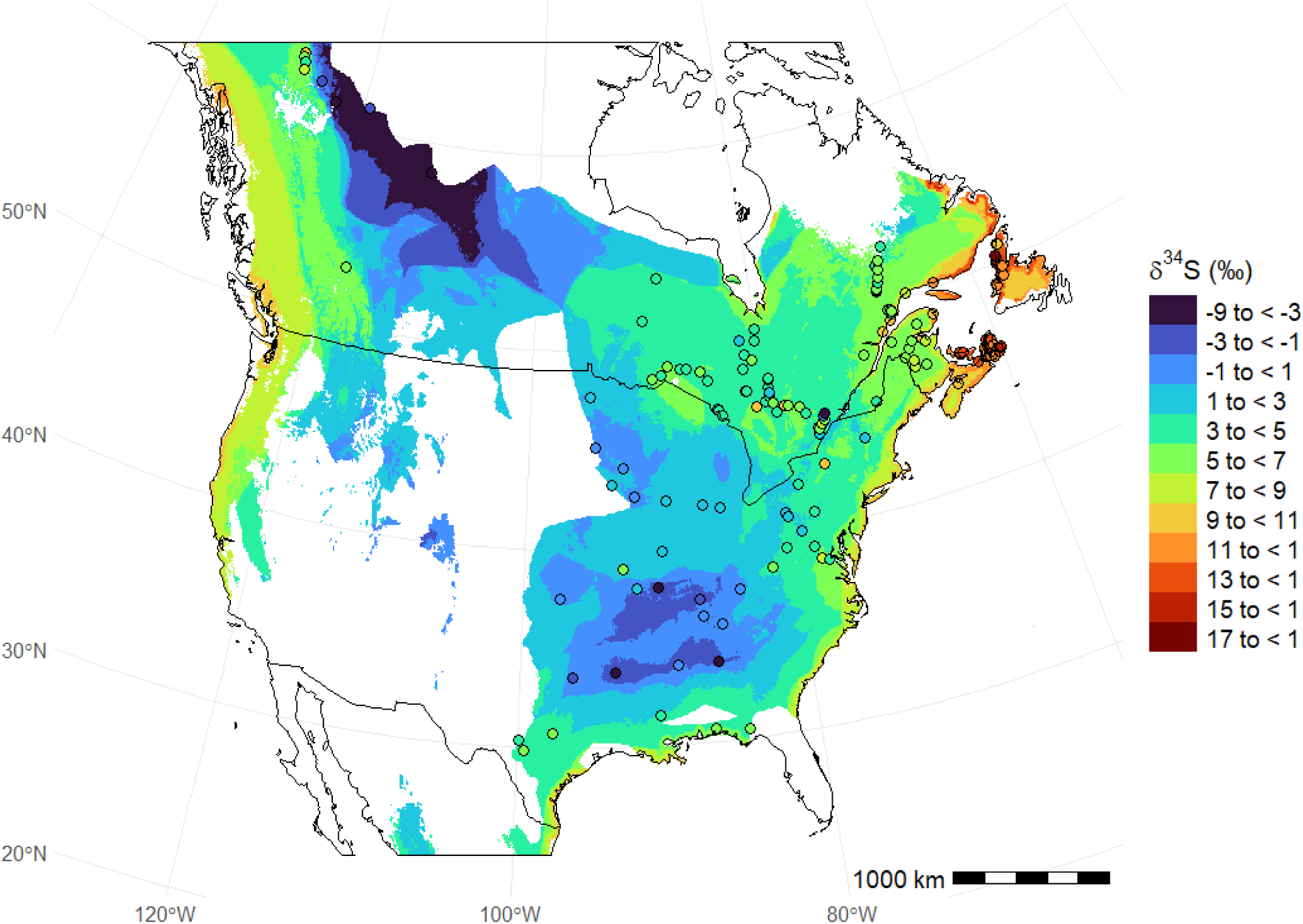
Predicted *δ*^34^S_foliar_ from random forest regression across eastern North America. Points represent plant sampling locations colored according to the *δ*^34^S_foliar_ color scale. Predictions are only provided for areas where predictors have values within the training set. Border polygons are from *rnaturalearth* (Massicotte and South 2023). See Supplementary Figure 3 for uncertainty layer.

#### 3.2. Relationship between plants and moths in the known-origin dataset

### 3.2.1 Calibration equation

We found a strong and significant relationship between *δ*^34^S_foliar_ and *δ*^34^S_moth_ values (F_1,73_=669.7, R^2^ = 0.90, p < 0.001, *δ*^34^S_moth_ = -0.08 + 0.94 *δ*^34^S_foliar_, see Figure 3), where moths have slightly lower o^34^S than the foliage they consume (i.e., -0.11 ‰ to 0.2 ‰ at the lower and higher ends of the foliar values we sampled, which is below our analytical error). An ANCOVA with known-origin o^34^S_moth_ value as the response variable, o^34^S_foliar_ value as covariate, and site nested within province as a factor showed that o^34^S_foliar_ was the strongest predictor (F_1,1_=1587, p < 0.001). Additionally, both province (F_1,1_=23, p < 0.001) and individual site within province (F_1,20_=5, p < 0.001), have minor but significant effects on o^34^S_moth_, possibly because these factors are correlated with local variation in *δ*^34^S values. Through the *calRaster* function of the *assignR* package, we obtained a strong correlation between *δ*^34^S_foliar_ isoscape and *δ*^34^S_moth_ values (*δ*^34^S_moth_ = -0.5 + 1.01 *δ*^34^S_foliar isoscape_, R^2^ = 0.94), and a similar calibration equation to the *δ*^34^S_foliar_-*δ*^34^S_moth_ regression. Differences between these two regressions are due to known-origin moths collected at five sites without available foliage. *δ*^34^S_foliar_ values are an adequate proxy to develop insect *δ*^34^S isoscapes.

**Figure 3:**
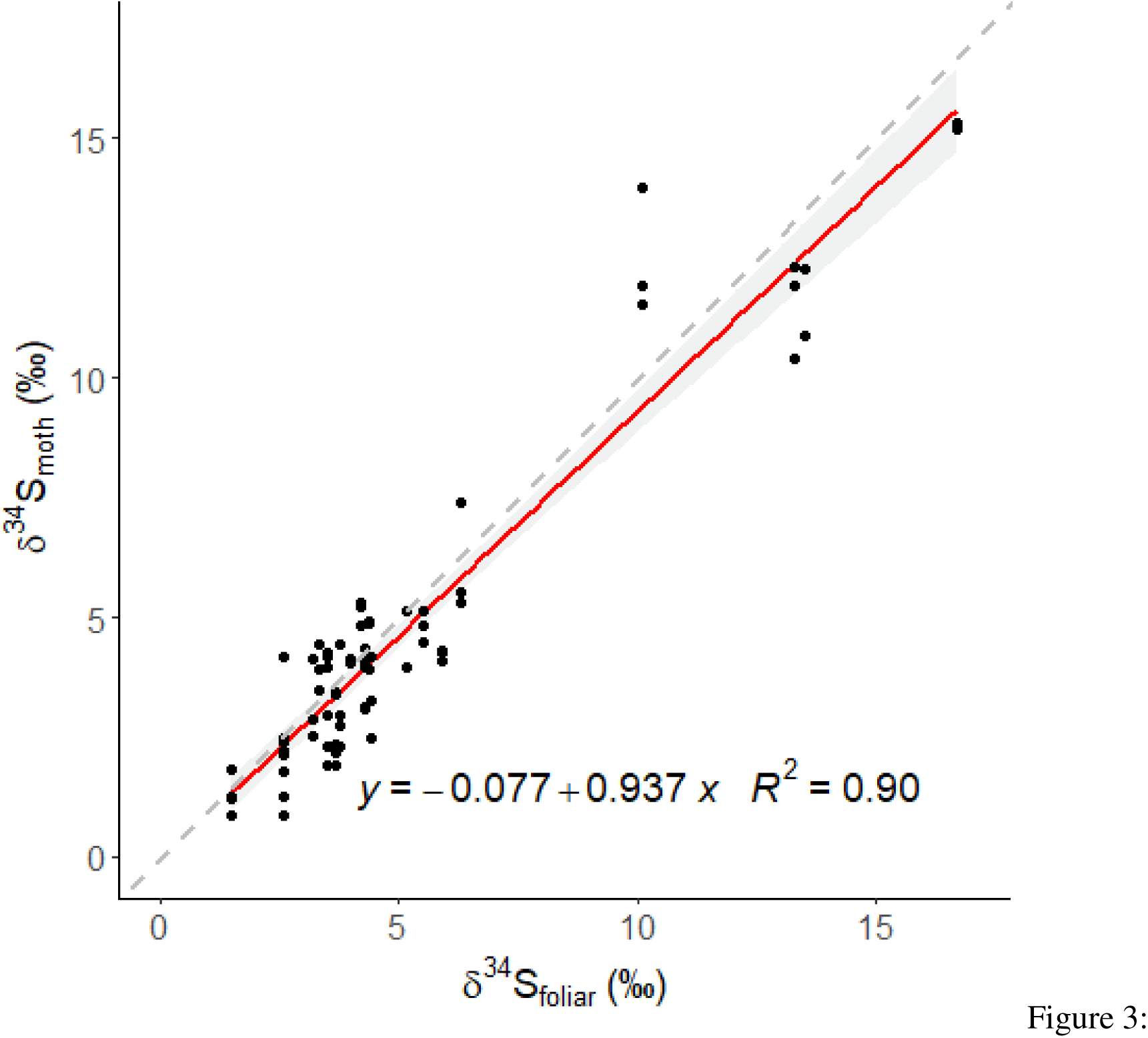
Relationship between individual moth *δ*^34^S values (*δ*^34^S_moth;_ n = 75) and the average tree *δ*^34^S value of their site of collection (*δ*^34^S_foliar;_ n = 23) was significant (F_1,73_= 669.7, p< 0.001) and explained 90% of the variance in *δ*^34^S_moth_ (*δ*^34^S_moth_ = -0.08 + 0.94\**δ*^34^S_foliar_). Grey band represents the 95% CI and broken line a 1:1 relationship between predicted and observed values.

### 3.2.2 Intra-site foliar *δ*^34^S variation

We observed limited *δ*^34^S_foliar_ intra-site variance associated with needle type and species. We found no difference between duplicates of the same needle tissue (21 independent trees). Duplicates showed an average *δ*^34^S_foliar_ difference of 0.16 ‰, well within the 0.3 ‰ analytical error. We found a small but nonetheless significant effect of foliage age on o^34^S values (paired *t-test*, p=0.021, t_6_=3.11), with young needles having a 0.78 ‰ higher value than old needles on the few sites studied (n=7). We found no significant effect of species when balsam fir and spruce *δ*^34^S_foliar_ were collected at the same site (paired *t-test*, p=0.83, t_8_=0.224). Intra-site *δ*^34^S_foliar_ variance was also similar between spruce (0.37 ‰, range: 0.12 - 0.85 ‰) and fir (0.56 ‰, range: 0.19 - 1.02 ‰), and both are similar to the analytical error.

### 3.2.3. Intra-site moths *δ*^34^S variations at known-origin sites

The average intra-site *δ*^34^S_moth_ standard deviation was 0.59 ‰ (range: 0.03 - 1.31 ‰). We used individual moth duplicates to test within moth variation, as a proxy for both within-sample natural variation in *δ*^34^S values and analytical error. We found no significant difference between samples and their duplicates (t_12_ = 0.48, p = 0.64), with a mean difference of 0.11 ‰.

#### 3.3 Quality assessment of isoscapes and geographic assignment

The *δ*^34^S_moth_ isoscape shows deviation from the expected proportion of samples correctly assigned (i.e., on the 1:1 line, Supplementary Figure 4A). This deviation, particularly at low probability quantile thresholds, is likely associated with overestimation of the error in the uncertainty raster. Overestimation of uncertainty has been reported when deriving uncertainty surfaces from quantile random forest (Bataille et al. 2022). *δ*^34^S data can be used to effectively eliminate large portions of the study area as probable natal origin (Supplementary Figure 4B), but could be further improved by refining the estimation of uncertainty. The odds ratio plot indicates that known-origin individuals are three times more likely to be assigned to their site of natal origin compared to random locations (Supplementary Figure 4C). These results demonstrate that sulfur isotopes are a viable method to trace the origin of insects, particularly for discriminating individuals from, or arriving at, coastal areas.

#### 3.4 Geographic assignments and intra-site variations of local individuals

As a pre-requisite to validate the application of *δ*^34^S geolocation, we generated geographic assignments of individuals captured in automated traps and classified as locals. 25 out of 32 individuals classified as locals through visual inspection of assignments (Arisaig n= 5 out of 6, Inverness n=3 out of 3, St. Modeste n= 3 out of 3, Baldwin n= 3 out of 3, Zinc Mine rd. n= 3 out of 7, Forestville n= 2 out of 2, Gaspe n= 3 out of 3, Pikauba n= 2 out of 3, Petit Etang n=1 out of 2) showed geographic assignments consistent with a possible local origin (Supplementary Figures 5-13) whereas the remaining individuals showed very low probability of origin at the capture site. Additionally, captured locals have a significantly higher intra-site *δ*^34^S_moth_ variance with an average standard deviation of 1.74 ‰ (range: 0.90 - 3.97 ‰) than those of known-origin sites (t_8.99_ = -3.33, p = 0.009) (Supplementary Figure 14).

3.5 Continuous-surface geographic assignment and atmospheric dispersal of immigrant moths Moths from Arisaig (n = 3) and Inverness (n = 3) categorized as immigrants in the automated traps exhibited a large degree of *δ*^34^S_moth_ variation and generally lower values than those predicted by the *δ*^34^S_moth_ isoscape for these coastal locations in Nova Scotia (Figure 4), confirming that they likely did not originate from the capture location. High probability regions included locations in northern New Brunswick, the Gaspé Peninsula and the north coast of the Saint Lawrence River (Figure 4, Supplementary Figure 15), and were highly distinct from those predicted for purported locals (Supplementary Figure 5, 6, 16). Among these areas, only some parts of the Gaspé Peninsula and the north shore of the Saint Lawrence, showed budworm defoliation in 2020, making these areas a more likely source of a mass migration event.

**Figure 4:**
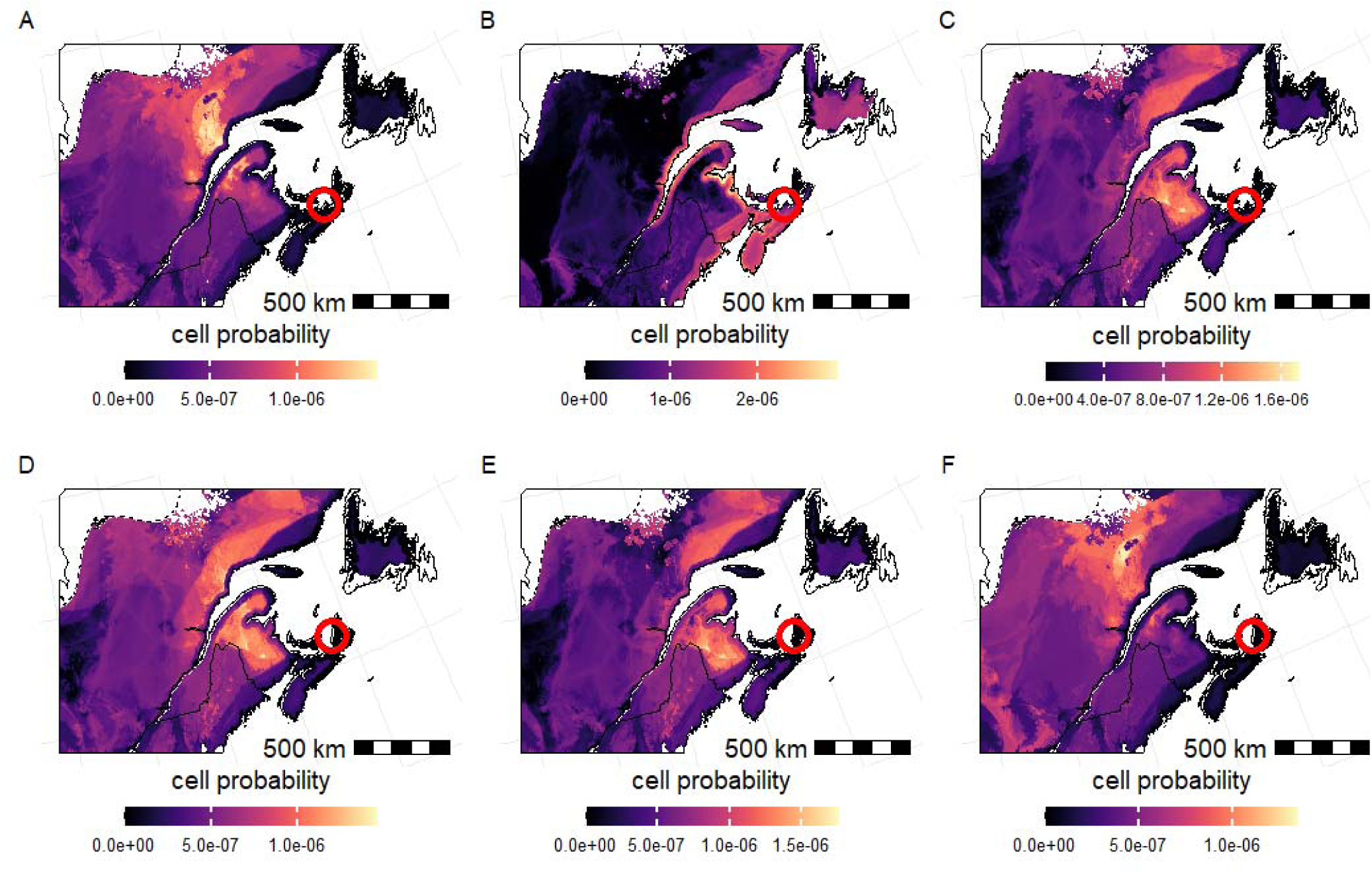
Posterior probability maps of all immigrant individuals sampled from Arisaig (A-C) and Inverness (D-F). Trap locations are shown as red circles. The bright yellow areas represent sites with a higher probability of origin; black areas represent zero probability. The probability values of each map sum to 1 and values are only provided for areas within the spruce budworm distribution range (broken line) where all isoscape predictors have values within the training set (Figure 2). Border polygons are from *rnaturalearth* (Massicotte and South 2023).

None of the simulated HYSPLIT trajectories within the approximate six-hour window of the capture period (i.e., 17:00h to 23:00h ADT) originated from areas of severe defoliation on the night of capture (July 22-23, 2020) indicating immigration either from earlier times and/or more distant areas. However, when reconstructing trajectories for the night before the capture event (July 21, 2020), 346 simulated trajectories originated from the outbreak zone on the southern part of the Gaspé Peninsula and landed less than 10km from the Arisaig and Inverness traps (Figure 5). Of these, all trajectories remained above an altitude of 250m, well above water and the tree canopy, whereas 11 trajectories were discarded because at one point before intersection the temperature fell below 15 °C (i.e., below the temperature flight threshold of spruce budworm moths) (Supplementary Figure 17). The temperature and altitude along the remaining trajectories (20.5 >°C ≥15; 600 m> altitude > 250 m) is consistent with the dispersal ecology of the moth, and would not have imposed physiological constraints or barriers to movement (Sanders et al. 1978). These remaining trajectories represent the most likely routes of the migrating eastern spruce budworm moths and include a water-crossing of over 400 km. The *δ*^34^S-based geographic assignment, defoliation maps, and successful HYSPLIT trajectories overlap in the southeastern Gaspé Peninsula, making this area the most likely natal origin of the moths that immigrated to Arisaig and Inverness on July 22-23, 2020

**Figure 5:**
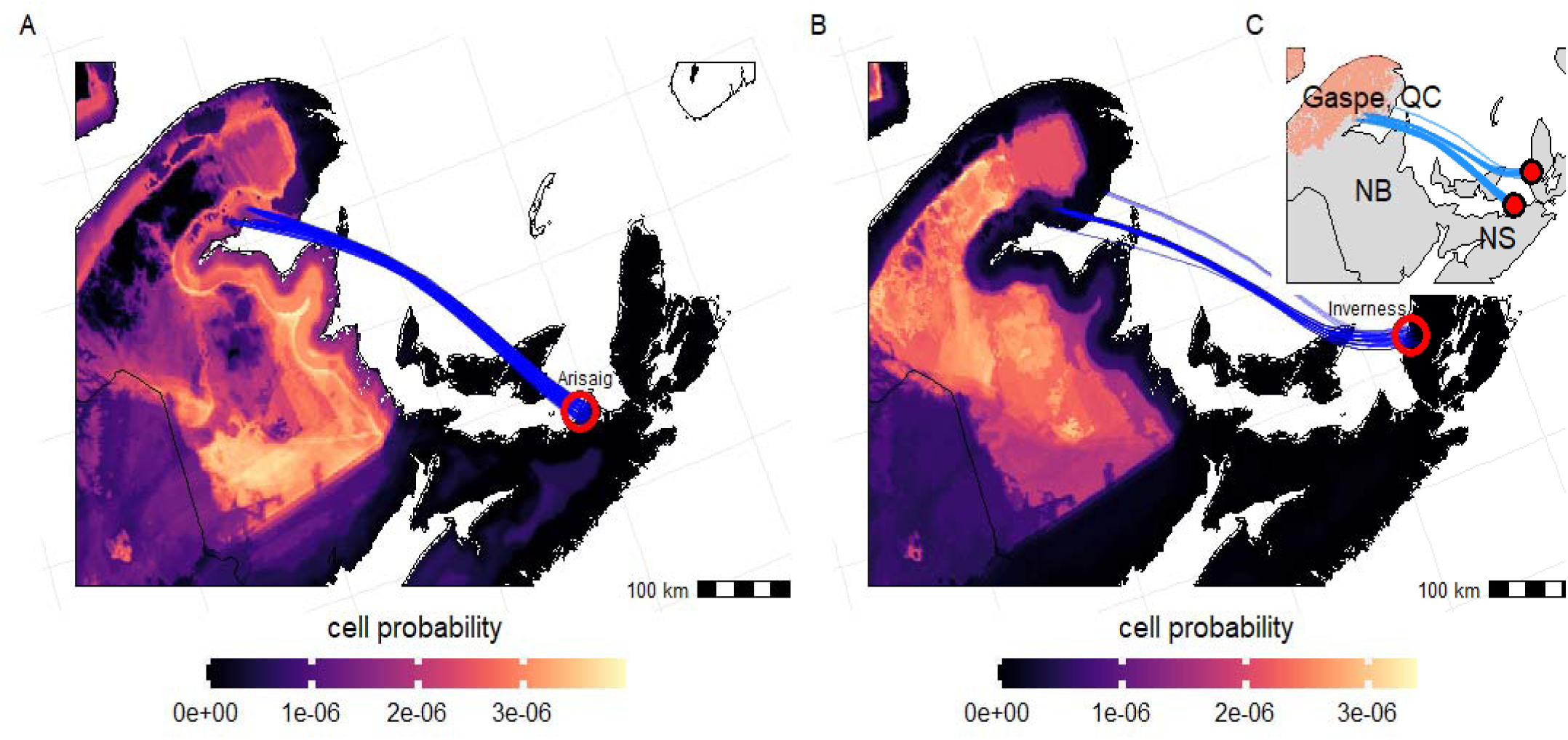
Joint posterior probability maps of all immigrant individuals sampled from Arisaig (A) and Inverness (B) with HYSPLIT trajectories (in blue) from 21 July 2020 departing from the defoliated zone and landing within 10 km from the automated traps. Trap locations are shown as red circles. Yellow areas represent sites of higher probability of origin; black areas represent zero probability of the individuals having originated there. The probability values of each map sum to 1 and values are only provided for areas within the spruce budworm distribution range (broken line) where all isoscape predictors have values within the training set (Figure 2). Border polygons are from *rnaturalearth* (Massicotte and South 2023). (C) Map depicting areas that were defoliated in 2020 (orange) and HYSPLIT trajectories (blue lines) to trap locations (red circles).

## 4. Discussion

### 4.1 Sulfur isoscape

The isoscape displays a progressive decline in *δ*^34^S_foliar_ values when moving away from the ocean (Figure 2), supporting previous studies about the dominance of marine sulfate deposition in controlling the continental *δ*^34^S variations in plants (Sparks et al. 2019), birds (Hebert and Wassenaar 2005, Brlík et al. 2023), and mammals (Zazzo et al. 2011, Bataille et al. 2021, Bataille et al. 2022). Partial dependence plots show that sea salt aerosol deposition rates, distance to the coast, wind, aridity index and potential evapotranspiration are important predictors of *δ*^34^S_foliar_ values. Interestingly, the relationship between *δ*^34^S_foliar_ and those predictors is not linear, with a sharp decline in the first few kilometers away from the coast, a plateau between 10-100 km from the coast (or when ecosystems become dryer and less windy), followed by a more gradual decline extending to more than 600 km (Figure 1), supporting previous findings on islands (Sparks et al. 2019).Wind and mean annual precipitation (represented by aridity index and potential evapotranspiration) display a plateau where wetter and windier areas have higher *δ*^34^S_foliar_ values, likely reflecting the typical wet and windy conditions within a 100 km band from the coast (Supplementary Figure 2). The low *δ*^34^S_foliar_ values away from the coast, likely also reflects the increasing contribution of terrestrial or anthropogenic sources with low *δ*^34^S in continental precipitation. These *δ*^34^S_foliar_ patterns are promising to discriminate within a gradient of coastal to continental migratory insects.

Deposition rates of mineral dust is also an important predictor of *δ*^34^S_foliar_, likely because mineral dust particles scavenge sulfates from the atmosphere and contribute to increase the sulfate deposition rate on ecosystems (Dupart et al. 2012). Consequently, areas with higher dust deposition have lower *δ*^34^S_foliar_ values because of the addition of terrestrial and anthropogenic sulfates with low *δ*^34^S to the ecosystems. For example, in the USA, the regions of high dust deposition surrounding the Midwest and southern USA, which have the highest rates of sulfur dioxide emissions, all show relatively low *δ*^34^S_foliar_ (Supplementary Figure 2). In summary, the *δ*^34^S_foliar_ isoscape reveals a predictable and broad range of spatial variation across our study area, which can be leveraged for geolocation, specifically to distinguish organisms that have metabolized local resources into their tissues at varying distances from the coast.

### 4.2 *δ*^34^S as a discrimination isotope for spruce budworm natal origins

The strong and nearly 1:1 relationship between *δ*^34^S_foliar_ and *δ*^34^S_moths_ indicates negligible or minimal fractionation of sulfur isotopes in this folivorous insect, supporting average findings for other animals (but note that some species deviate from this average; Raoult et al. 2024).

Additionally, the limited variability between plant species and age also supports relatively homogeneous *δ*^34^S values within ecosystems. This conservation of *δ*^34^S values within tissues, species and across trophic levels highlights the dominance of spatial processes in controlling sulfur isotope patterns, and makes sulfur isotopes attractive for isotope geolocation relative to other elements that are strongly influenced by trophic and metabolic processes (e.g., hydrogen, carbon, or nitrogen).

We found low but non-negligible amounts of variation among individual known-origin moths from the same site (median standard deviation = 0.59 ‰), whereas putative local moths showed significantly higher *δ*^34^S_moth_ variations (mean standard deviation = 1.74 ‰), particularly at sites close to the ocean. This high intra-site *δ*^34^S variability for purported local moths mirrors the high variability of hydrogen and strontium isotopes at those sites, further supporting significant local mobility of spruce budworm moths beyond what had been reported in the literature before the use of isotope tracers (Dargent et al. 2023). These observations suggest that “local” moths could move well beyond the few hundred meters reported in early studies by Sanders (1983) and/or that existing qualitative classification based on time of capture are not always effective at identifying migrants. The extent of this local movement ought to be explored further as it informs the extent to which outbreaks can expand passively.

### 4.3 A new framework to trace insect pests

Single-tool approaches to tracing pest movement have limitations, but by combining different and independent approaches we can reduce those constraints and provide unprecedented detailed analysis of pest immigration events. Arisaig and Inverness immigrants captured on the night of July 23^rd^ 2020 showed areas of high probability of natal origin around the Gaspé Peninsula, the northern shore of the Saint Lawrence, and New Brunswick (Figure 5). The *δ*^34^S-derived estimates of natal origin exclude large parts of the study area. Some of those regions of high probability coincide with regions that were defoliated in 2020, making them strong candidate for dispersal source-areas supported by independent ecological evidence associated to spruce budworm migratory patterns (i.e., the budworm density-defoliation patterns, and population density-emigration behaviour, Régnière and Nealis 2019). Interestingly, the most probable areas of origin obtained from wind dispersal trajectories coincided with areas of high probability of natal origin based on *δ*^34^S_moth_ values. Additionally, the absence of local moth captures at Arisaig and Inverness on the days preceding mass capture, combined with the lack of evidence for high larvae densities indicative of high population levels in the vicinity of the traps (nor even in New Brunswick), further support the non-local origin of these individuals. The most parsimonious explanation to integrate these results is that those immigrant moths found in Arisaig and Inverness dispersed on the previous night (i.e., July 21^st^) from the Gaspe Peninsula. Together, isotope geolocation, defoliation maps, and atmospheric dispersal models, along with the network of automated pheromone traps with sensors to determine capture times, provides an unprecedented and high spatiotemporal resolution view of the ecology and migration characteristics of this major insect pest.

In this case study, we demonstrated that *δ*^34^S values are particularly useful to identify and track the migration of insects originating from areas located at different distances from the coast, where we see the steepest *δ*^34^S variations. However, the accuracy of these geographic assignments could be further improved by more extensive sampling of the study region, particularly in locations with *δ*^34^S values between 7‰ and 12‰ (Figure 1A), and improvements to estimating spatial uncertainty of the isoscape using quantile random forest, which tends to overestimate uncertainty (Bataille et al. 2022). While the precision of the *δ*^34^S geolocator will be strongest to track insect migration in this type of environmental setting, a key strength of isotope geolocation is the possibility to combine multiple isotopes. Using hydrogen and strontium isotopes along with sulfur, can considerably increase the precision of geolocation (Bataille et al. 2021, Dargent et al. 2023), enabling the tracking of insect pests at fine spatiotemporal resolution in a range of environmental conditions.

### 4.4. Management implications

Our new integrated tracking framework advances the possibility of sustainably managing insect pests. Knowing whether immigration events occur at a sub-regional (e.g., tens of kilometres) or regional (e.g. hundreds of kilometres) scale can help assess the required scale of coordination between actors involved in pest management. For example, for spruce budworm, sub-regional immigrations are more likely to concern provincial or local jurisdiction management strategies, whereas regional, long-distance, immigration would require higher cooperation among provinces to effectively manage outbreaks. So far, Canadian government scientists have informed management of spruce budworm moth outbreaks through the Spruce Budworm Early Intervention Strategy (e.g., Johns et al. 2019) which uses BTK (i.e., a bacteria-derived toxin that targets Lepidoptera larvae) to spray areas of rapidly growing populations and keep them below outbreak levels. Our study demonstrates that eastern spruce budworm moths can migrate hundreds of kilometers across provincial borders. Thus, spruce budworm outbreak management would benefit from inter-provincial scale cooperation and coordination, as outbreaks in one province can spread to others through migration.

The integrated framework we propose is transferable to evaluate, at high temporal and spatial resolution, the migration ecology of many forest and agricultural insect pests, as well as migratory insects of conservation interest. Combining automated pheromone traps with isotopic analyses and atmospheric dispersal models enables the precise identification of immigrants and their origins, and offers the possibility to study the biotic, climatic, meteorological and environmental controls of outbreak spread. Such knowledge could be leveraged along with high spatiotemporal resolution remote-sensing of population data to predict in nearly real-time the population dynamics and outbreak spread risks of a given insect pest. Using these quantitative monitoring and forecasting tools, researchers could considerably improve the management or conservation strategies of migratory insects in both present and future climate scenarios.

## 5. Conclusion

Our study introduces a novel, integrated framework that combines automated traps equipped with sensors, isotopic analysis, specifically *δ*^34^S geolocation, field surveys of defoliation representative of ecological dynamics (i.e., population density and mass exodus triggers), and atmospheric dispersal modeling with physiological constraints (i.e., inclusion of insect flight velocity, temperature limits on flight, and altitude) to effectively trace the migration of insect pests. This framework provides the possibility to track the origin and routes of insect migration at high spatio-temporal resolution, information that is crucial for understanding the ecological conditions triggering outbreak spread. By enabling precise identification of immigrant origins and routes, this novel framework provides crucial data for enhancing cooperation between management and stakeholders and for making management strategies targeted, scalable and sustainable. While this framework was demonstrated on spruce budworm, it is entirely transferable and promises significant advancements in the sustainable management of various forest and agricultural insect pests. The combination of automated trap monitoring with AI identification, wind trajectory modeling, radar, high-resolution remote sensing, and next-generation isotope geolocation, goes beyond insect pest management and also applies to other migratory insects of conservation or bio-surveillance interest. Developing such integrated monitoring networks across the globe is a promising avenue to evaluate and manage the global risks and benefits of migratory insects for ecosystems, food production, and human societies in the face of global change.

## Acknowledgements

Several colleagues helped with collections, in particular, CFS: J. Bowden, S. Bourassa, M. Gauvin, W. MacKinnon, E. Moise, D. Pureswaran, J. Warren; Provincial Governments: NL: T. Rideout, B. Brake, B. Hinks, J. Motty, NS: J. Ogden, A. McGill, QC: J-J. Bertrand, P. Therrien; Parks Canada: S. Arnold, H. Lightfoot; Invasive Species Centre: D. Sparks, D. Dutkiewicz. A. Vanio Matila, M Lemire helped with sample preparation. Methods for isotope analyses were developed by the team at the Jan Veizer Isotope Laboratory (U Ottawa).

## Author contributions

FD, J-NC, and CPB: conceived and designed the project. FD, J-NC, KP, and MR: collected data. FD, MM, NB, JA, prepared and analysed samples. FD, J-NC, KS, MR and CPB: analyzed the data. FD, J-NC and CPB: contributed reagents/materials/analysis tools. FD: wrote the first draft and revised it with input from coauthors. FD, J-NC, CPB: revised and edited the manuscript. All authors reviewed and contributed to the article, and approved the submitted version.

