## Supplement for "A novel integrated framework to identify and characterize regional-scale pest insect dispersal"

Supplementary Materials

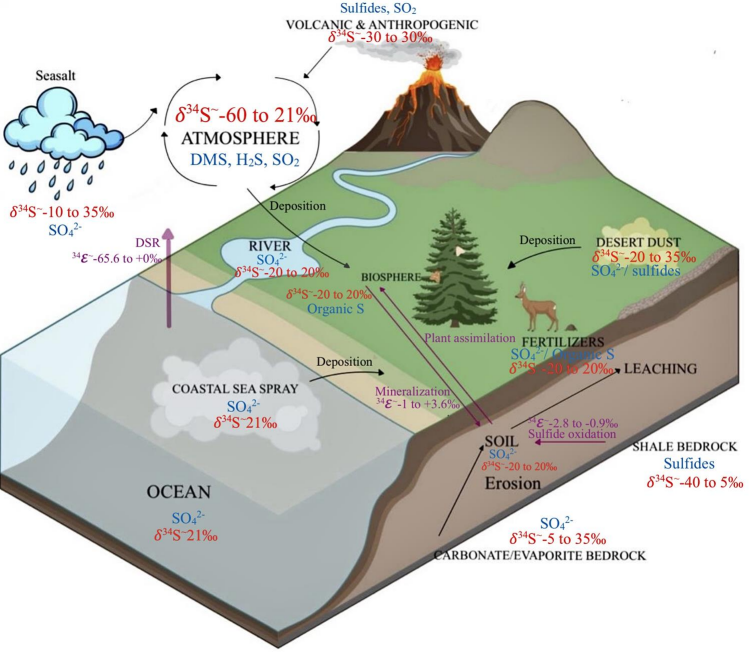
Supplementary Figure 1: Overview of the sulfur cycle and expected *δ*^34^S values in different environmental components

Sulfur isotopes (referred to as *δ*^34^S) vary on the landscape with atmospheric deposition processes or geology and have shown promising potential for geolocation in archeology (Clément P. Bataille et al., 2021) and bird ecology (Brlík et al., 2024; Brlík et al., 2023). Newton (2021) demonstrated that moths collected across the UK showed consistent spatial patterns with *δ*^34^S gradient from the coast to inland. However, *δ*^34^S has not been tested as a geolocator for insect migration. Sulfur isotopes are attractive as an insect geolocator because sulfur is a macronutrient present in measurable amounts in many insect tissues through two protein-forming amino acids, the essential methionine and the non-essential cysteine (Tcherkez & Tea, 2013). The sulfur isotope cycle is also well-characterized, primary producers (i.e., plants) obtain sulfur primarily from soils and more rarely through atmospheric uptake as sulfur oxides and carbonyl sulfides (Tcherkez & Tea, 2013; Trust & Fry, 1992). In most ecosystems, soil sulfur is derived from atmospheric deposition of marine sulfates deposited either by short-distance dry deposition of sea spray or wet deposition of sulfates dissolved in precipitation (Figure 1). The modern ocean sulfates have homogeneous and elevated *δ*^34^S +21 ± 0.2 ‰ (Böttcher, Brumsack, & Dürselen, 2007). Coastal ecosystems with high rates of marine sulfate inputs have usually higher *δ*^34^S values than those located more inland (Clément P. Bataille et al., 2021; Sparks, Crowley, Rutherford, & Jaggernauth, 2019). Conversely, in ecosystems with low marine sulfate deposition, other processes often dominate sulfur cycling including sulfates from sulfide oxidation from soil minerals (e.g., black shales) with very low *δ*^34^S values (<0 ‰), weathering of sulfate-rich rocks (e.g., evaporites) with high but variable *δ*^34^S values, atmospheric deposition of sulfates from human emissions with generally low *δ*^34^S values or dust deposition of sulfur-containing minerals from desertic areas or volcanic ash with lower *δ*^34^S values (Nehlich, 2015), and sulfide accumulation from bacteria and archaea activities in under thawing permafrost with extremely low *δ*^34^S values (Harrison & Thode, 1958; Stevens et al., 2023). The complex cycling and multiple isotopically-distinct sources of sulfur in ecosystems might lead to isotope patterns independent from hydrogen and strontium providing a new tool for high-resolution insect tracing.

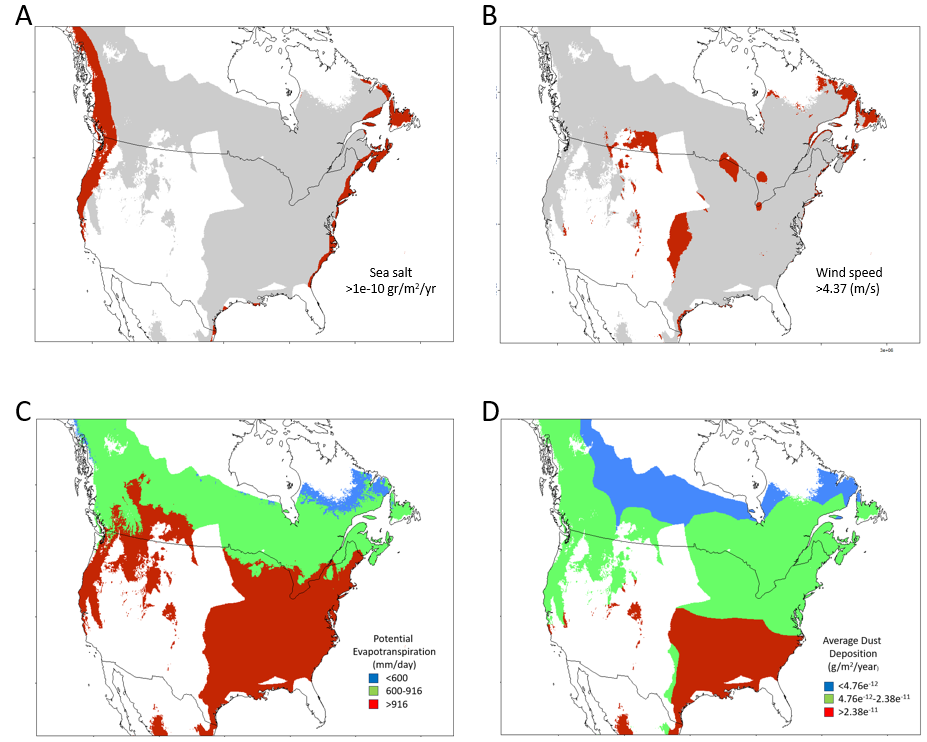

Supplementary Figure 2: Evaluation of spatial thresholds based on partial dependence plots. (A) sea salt aerosol deposition (g/m^2^/yr), (B) wind speed (m/s), (C) potential evapotranspiration (mm/day), and and (D) mineral dust deposition (g/m^2^/yr). Values are only provided for areas where all predictors have values within the training set. Border polygons are from *rnaturalearth* (Massicotte & South, 2023)

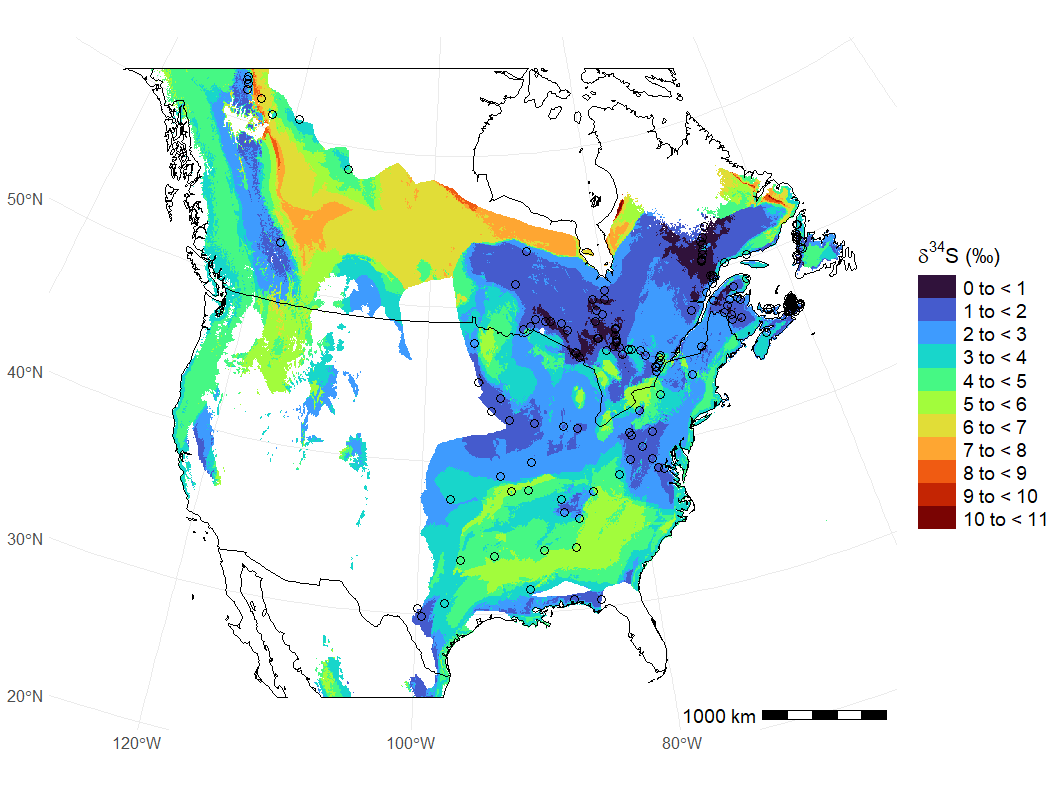

Supplementary Figure 3: Spatially-explicit uncertainty from the random forest predictions. The uncertainty represents one standard deviation and was obtained from quantile random forest modeling (see Methods). Empty circles indicate locations where foliar samples were collected. As for Figure 2, predictions are only provided for areas where predictors have values within the training set. Border polygons are from *rnaturalearth* (Massicotte & South, 2023).

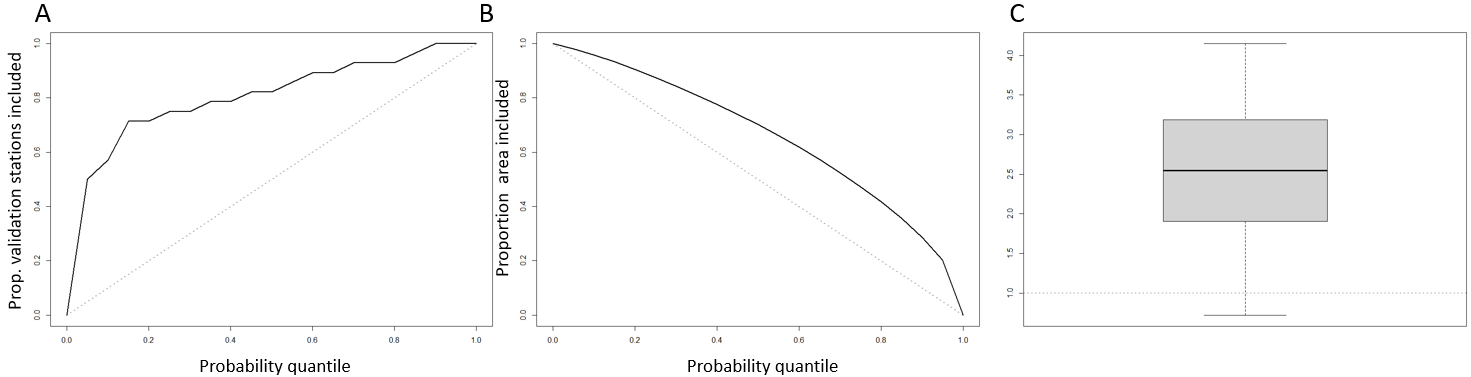

Supplementary Figure 4: Quality assessment plot of *δ*^34^S_moth_ isoscape using known-origin spruce budworm moth. The *QA* function in *calRaster* (Ma et al. 2020) randomly splits the known-origin data into calibration and validation subsets, recalibrates the foliar isoscape using the calibration set and assigns samples from the validation set, repeating this iteratively. (A) A measure of bias, as the proportion of validation samples that are correctly assigned based on the probability threshold. When accurate, the posterior probabilities fall along the 1:1line, higher values suggest overfitting. (B) A measure of granularity which shows the proportion of study area (Eastern Canada) that is excluded from assignments of origin based of the probability threshold. A higher curve denotes capacity for more precise assignments. (C) The posterior probability of all the known-origin locations relative random locations, where higher values indicate a higher odd of an individual being assigned to its known location than a random one, akin to stronger support for a given site relative to others.

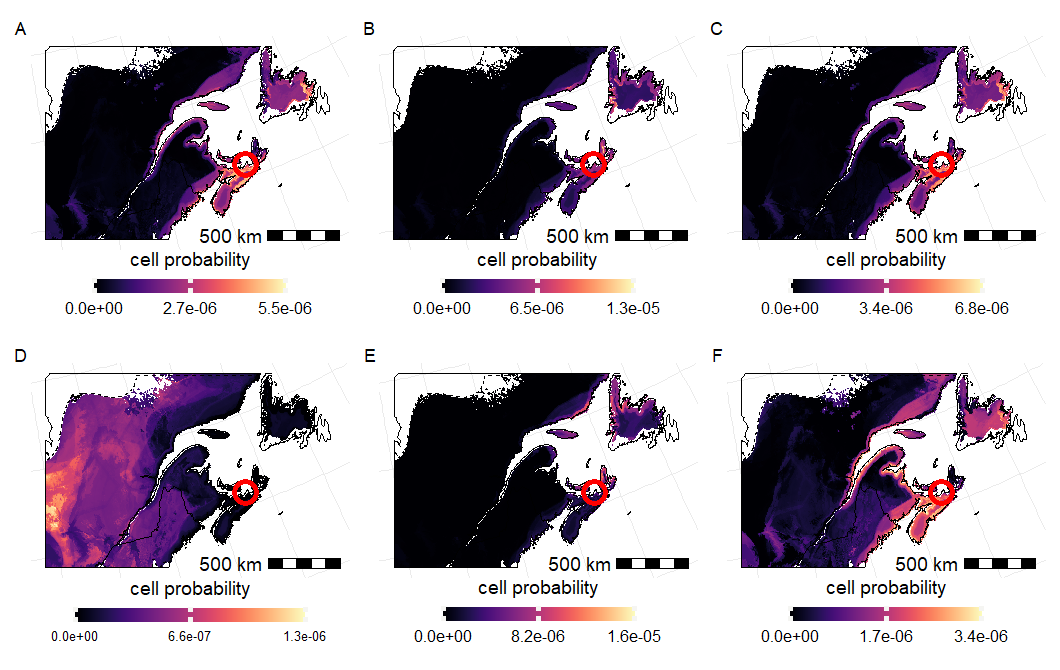

Supplementary Figure 5: Posterior probability maps of all six individuals sampled from the Arisaig (NS) trap that were classified as local based on capture time inferences (A-F). Bright yellow areas represent sites of higher probability of origin, black areas represent no probability of the tissue having originated there. Trap location shown as red circle. All cells in model sum up to a probability of 1. Values are only provided for areas where all predictors have values within the training set and within the spruce budworm distribution range (broken line). Border polygons are from *rnaturalearth* (Massicotte & South, 2023). Five of the individuals have a *δ*^34^S value similar to that of their capture area, confirming the classification of local (A-C, E-F). However, one individual had a *δ*^34^S value that suggests that it originated from a non-local area (D).

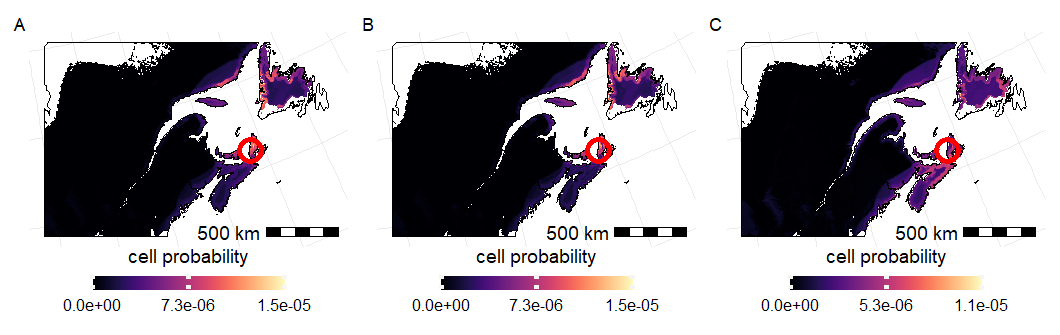

Supplementary Figure 6: Posterior probability maps of all three individuals from the Inverness (NS) trap that were classified as local based on capture time inferences (A-C). Bright yellow areas represent sites of higher probability of origin, black areas represent no probability of the tissue having originated there. Trap location shown as red circle. All cells in model sum up to a probability of 1. Values are only provided for areas where all predictors have values within the training set and within the spruce budworm distribution range (broken line). Border polygons are from *rnaturalearth* (Massicotte & South, 2023). All individuals have a *δ*^34^S value similar to that of their capture area, confirming the classification of local.

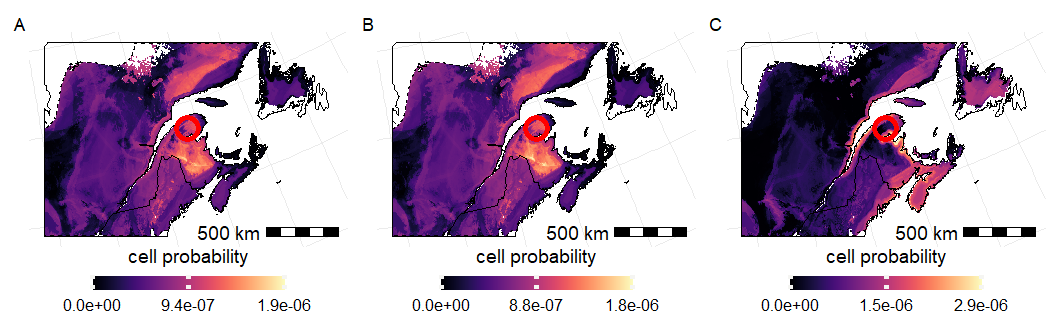

Supplementary Figure 7: Posterior probability maps of all three individuals from the Baldwin (QC) trap that were classified as local based on capture time inferences (A-C). Bright yellow areas represent sites of higher probability of origin, black areas represent no probability of the tissue having originated there. Trap location shown as red circle. All cells in model sum up to a probability of 1. Values are only provided for areas where all predictors have values within the training set and within the spruce budworm distribution range (broken line). Border polygons are from *rnaturalearth* (Massicotte & South, 2023). All of the individuals have a *δ*^34^S value similar to that of their capture area, confirming the classification of local.

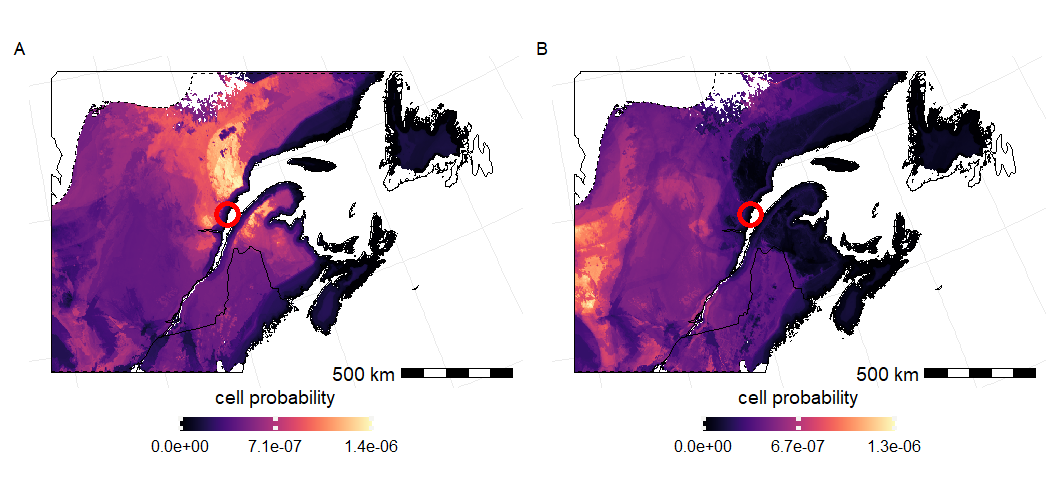

Supplementary Figure 8: Posterior probability maps of all two individuals the Forestville (QC) trap that were classified as local based on capture time inferences (A-B). Bright yellow areas represent sites of higher probability of origin, black areas represent no probability of the tissue having originated there. Trap location shown as red circle. All cells in model sum up to a probability of 1. Values are only provided for areas where all predictors have values within the training set and within the spruce budworm distribution range (broken line). Border polygons are from *rnaturalearth* (Massicotte & South, 2023). Individuals A and B, both have values that could originate in their capture area, but with low probability.

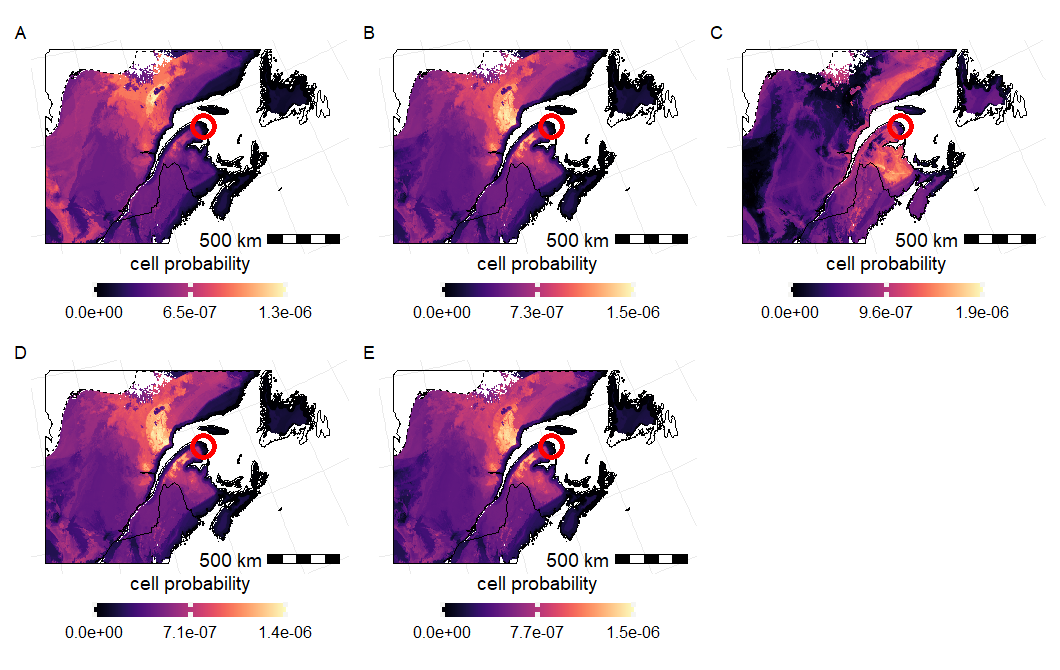

Supplementary Figure 9: Posterior probability maps of all three individuals from the Gaspe (QC) trap that were classified as local based on capture time inferences (A-C), with a duplicate of individual A using the same tissues (i.e. head and thorax) (D), and another duplicate of A using abdomen tissue (E). Bright yellow represent sites of higher probability of origin, clack areas represent no probability of the tissue having originated there. Trap location shown as red circle. All cells in model sum up to a probability of 1. Values are only provided for areas where all predictors have values within the training set and within the spruce budworm distribution range (broken line). Border polygons are from *rnaturalearth* (Massicotte & South, 2023). The three individuals (A-C), and the replicates of individual A (D-E), have a *δ*^34^S value similar to that of their capture area, but with low probability of origin.

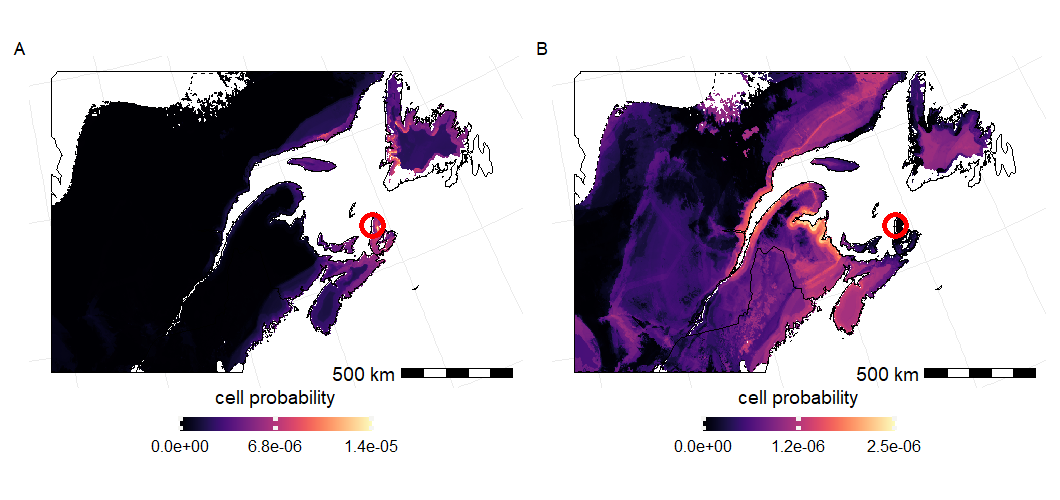

Supplementary Figure 10: Posterior probability maps of all individuals from the Petit Étang (NS) trap that were classified as local based on capture time inferences (A-B). Bright yellow areas represent sites of higher probability of origin, black areas represent no probability of the tissue having originated there. Trap location shown as red circle. All cells in model sum up to a probability of 1. Values are only provided for areas where all predictors have values within the training set and within the spruce budworm distribution range (broken line). Border polygons are from *rnaturalearth* (Massicotte & South, 2023). Individual A had a *δ*^34^S value similar to that of its capture area, confirming the classification of local, whereas individual B had a *δ*^34^S value that suggests that it originated from a non-local area.

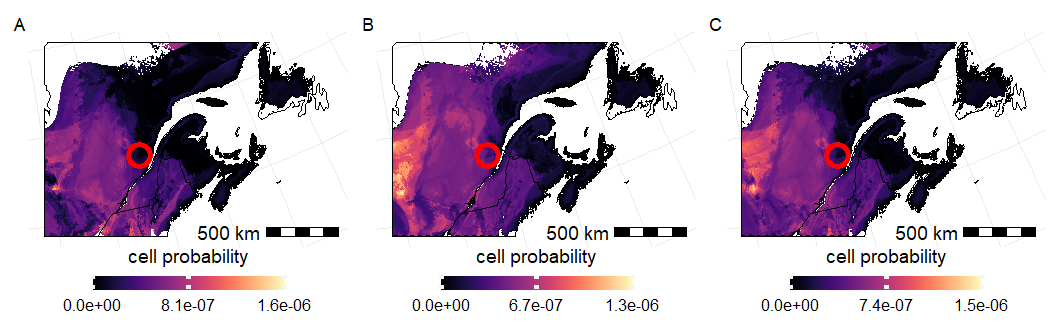

Supplementary Figure 11: Posterior probability maps of all three individuals from the Pikauba (QC) trap that were classified as local based on capture time inferences (A-C). Bright yellow areas represent sites of higher probability of origin, black areas represent no probability of the tissue having originated there. Trap location shown as red circle. All cells in model sum up to a probability of 1. Values are only provided for areas where all predictors have values within the training set and within the spruce budworm distribution range (broken line). Border polygons are from *rnaturalearth* (Massicotte & South, 2023). Two of the individuals have a *δ*^34^S value similar to that of their capture area, confirming the classification of local (B-C). However, one individual had a *δ*^34^S value that suggests that it originated from a non-local area (A).

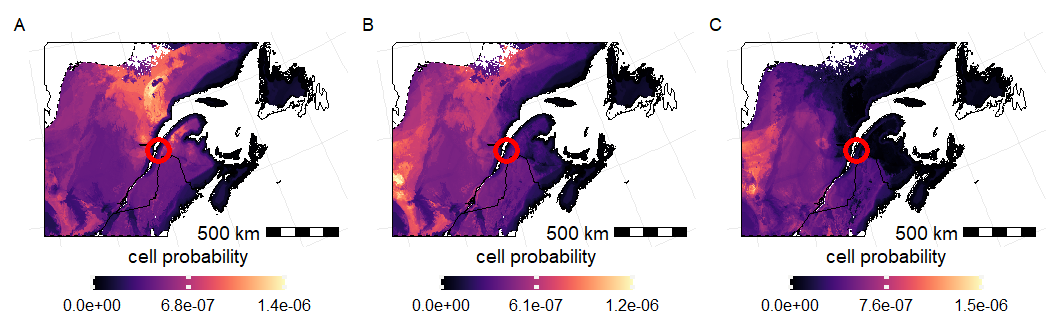

Supplementary Figure 12: Posterior probability maps of all three individuals from the Sainte Modeste (QC) trap that were classified as local based on capture time inferences (A-C). Bright yellow areas represent sites of higher probability of origin, black areas represent no probability of the tissue having originated there. Trap location shown as red circle. All cells in model sum up to a probability of 1. Values are only provided for areas where all predictors have values within the training set and within the spruce budworm distribution range (broken line). Border polygons are from *rnaturalearth* (Massicotte & South, 2023). All of the individuals have a *δ*^34^S value similar to that of their capture area, confirming the classification of local, but one of those individuals has a low probability of origin (C).

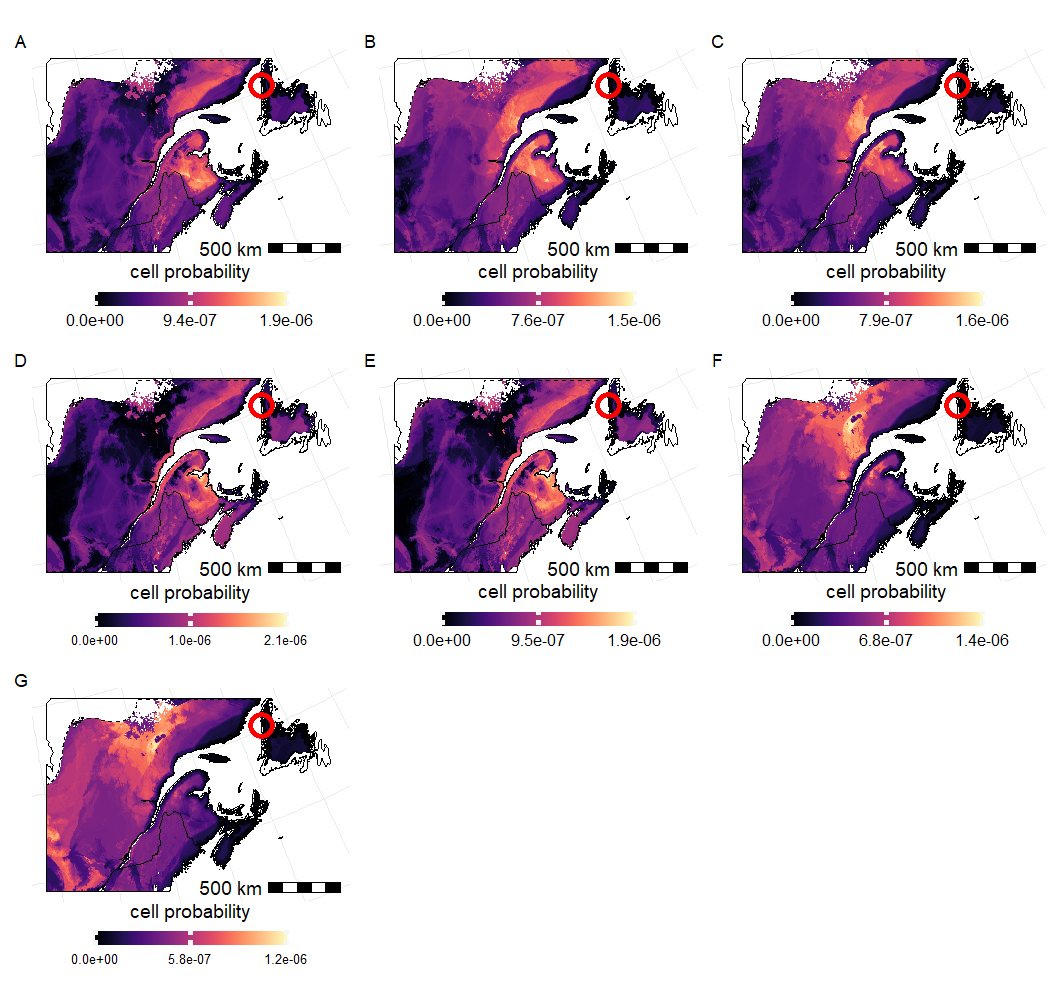

Supplementary Figure 13: Posterior probability maps of all seven individuals from the Zinc Mine road (NL) trap that were classified as local based on capture time inferences (A-G). Bright yellow areas represent sites of higher probability of origin, black areas represent no probability of the tissue having originated there. Trap location shown as red circle. All cells in model sum up to a probability of 1. Values are only provided for areas where all predictors have values within the training set and within the spruce budworm distribution range (broken line). Border polygons are from *rnaturalearth* (Massicotte & South, 2023). Most individuals had a *δ*^34^S value that suggests that it originated from a non-local area (B-C, F-G). However, three individuals have a *δ*^34^S value similar to that of their capture area, albeit with low probability (A, D-E).

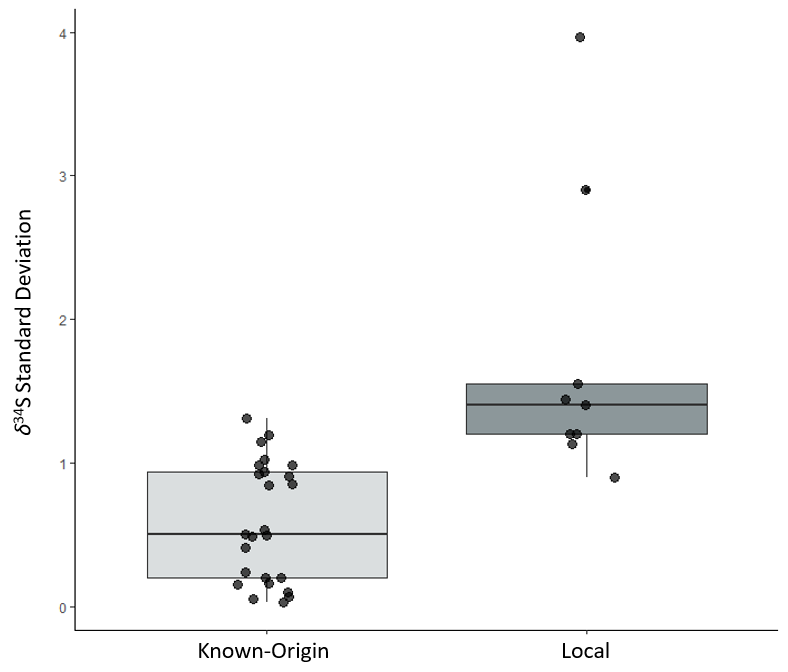

Supplementary Figure 14: Box and whiskers plot of known origin (i.e., moths collected as pupae) and putative locals (i.e., inferred by time and date of capture on automated traps) within-site *δ*^34^S_moth_ variation. Each point represents the within-site standard deviation in *δ*^34^S_moth_ value among individuals. Mean standard deviation (known-origin: 0.59 ‰, locals: 1.74‰) was significantly different between groups (t_8.99_ = -3.33, p = 0.009).

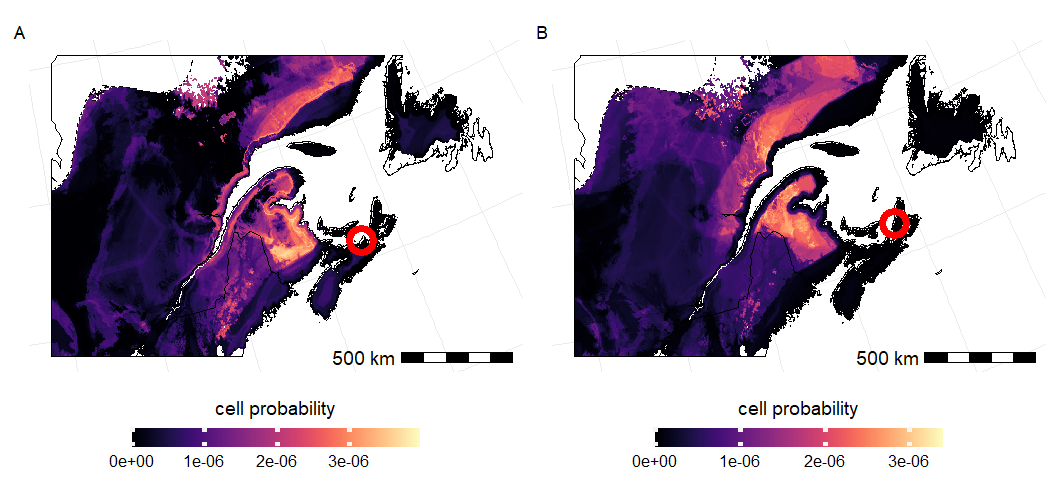

Supplementary Figure 15: Joint posterior probability of origin map for all immigrants sampled from Arisaig (A, n=3) and Inverness (B, n=3). Joint probability corresponds to the probability of all three individuals coming from each grid cell in the analysis area, bright yellow values indicate higher probability of origin in a given cell, whereas black values indicate no probability of origin. Trap location shown as red circle. All cells in model sum up to a probability of 1. Values are only provided for areas where all predictors have values within the training set and within the spruce budworm distribution range (broken line). Border polygons are from *rnaturalearth* (Massicotte & South, 2023).

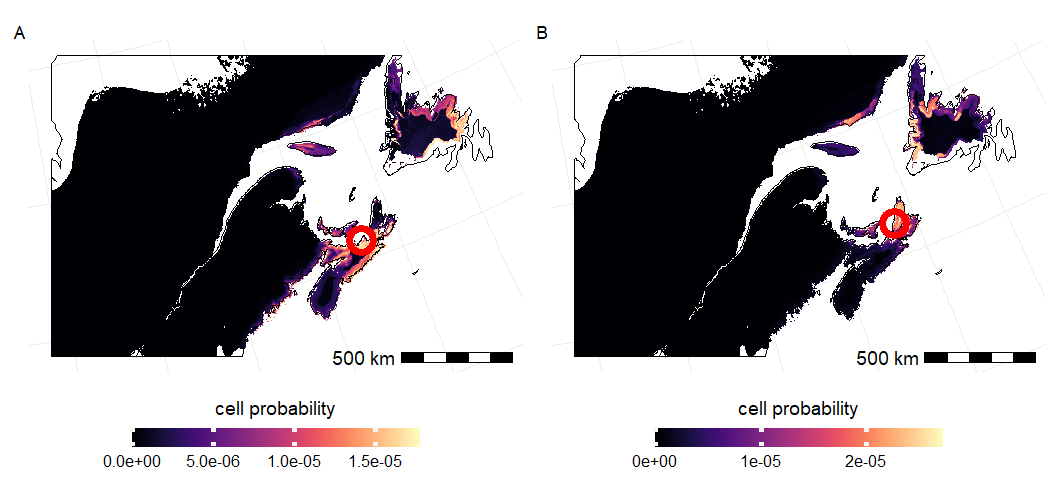

Supplementary Figure 16: Joint posterior probability of origin map for five out of six locals captured at Arisaig (A) and all three locals capture at Inverness (B). Joint probability corresponds to the probability of all individuals coming from each grid cell in the analysis area, bright yellow values indicate higher probability of origin in a given cell, whereas black values indicate no probability of origin. Trap location shown as red circle. All cells in model sum up to a probability of 1. Values are only provided for areas where all predictors have values within the training set and within the spruce budworm distribution range (broken line). Border polygons are from *rnaturalearth* (Massicotte & South, 2023).

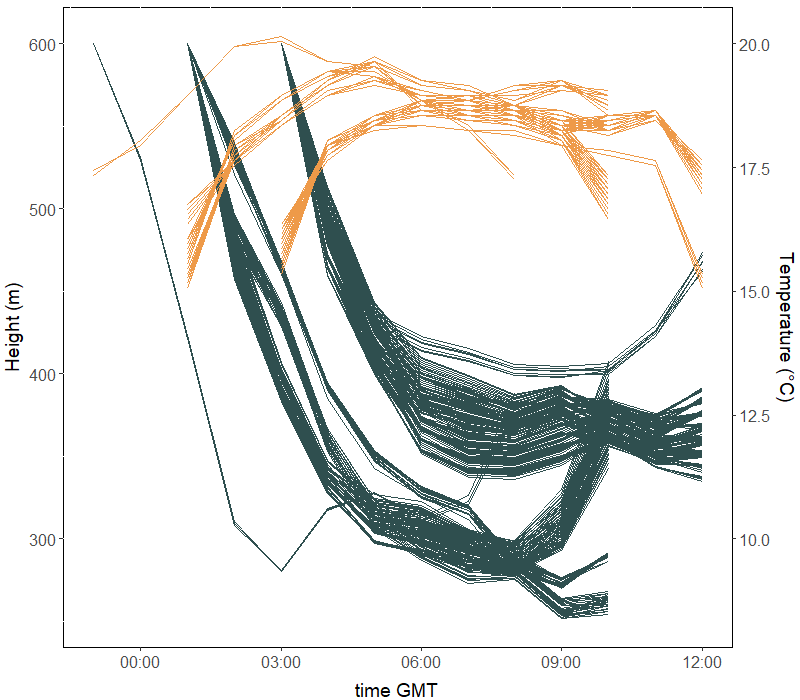

Supplementary Figure 17: Altitude and temperature along the trajectories simulated using HYSPLIT. Lines represent each individual trajectory for altitude (green, left axis) and temperature (orange, right axis). Time is on GMT, simulations were started at equivalent 19 h, 21 h, and 23 h eastern time, and allowed to continue for nine hours.
